# Fast Estimation of Recombination Rates Using Topological Data Analysis

**DOI:** 10.1101/395210

**Authors:** Devon P. Humphreys, Melissa R. McGuirl, Michael Miyagi, Andrew J. Blumberg

## Abstract

Accurate estimation of recombination rates is critical for studying the origins and maintenance of genetic diversity. Because the inference of recombination rates under a full evolutionary model is computationally expensive, an alternative approach using topological data analysis (TDA) has been proposed. Previous TDA methods used information contained solely in the first Betti number (*β*_1_)of the cloud of genomes, which relates to the number of loops that can be detected within a genealogy. While these methods are considerably less computationally intensive than current biological model-based methods, these explorations have proven difficult to connect to the theory of the underlying biological process of recombination, and consequently have unpredictable behavior under different perturbations of the data. We introduce a new topological feature with a natural connection to coalescent models, which we call *ψ*. We show that *ψ* and *β*_1_ are differentially affected by changes to the structure of the data and use them in conjunction to provide a robust, efficient, and accurate estimator of recombination rates, TREE. Compared to previous TDA methods, TREE more closely approximates of the results of commonly used model-based methods. These characteristics make TREE well suited as a first-pass estimator of recombination rate heterogeneity or hotspots throughout the genome. In addition, we present novel arguments relating *β*_1_ to population genetic models; our work justifies the use of topological statistics as summaries of distributions of genome sequences and describes a new, unintuitive relationship between topological summaries of distance and the footprint of recombination on genome sequences.

---

Recombination is a fundamental source of genetic variation in many natural populations. By bringing existing mutations into novel genomic backgrounds, recombination can accelerate the rate at which adaptation occurs, as well as prevent the buildup of deleterious variants which occurs in asexuals via Muller’s ratchet (1–3). It is therefore critical to measure the rates of recombination in order to understand rates of adaptation. Resolution along the genome is also an important factor, as recombination rates are known to vary substantially along chromosomes. In particular, hotspots of recombination have been found associated with a variety of sequence and structural motifs in natural populations (4–8). In addition to hotspot detection, better estimation techniques for recombination rates can also improve our understanding of observed levels of linkage disequilibrium in genome data (9), and consequently the expected signatures of various evolutionary phenomena such as selective sweeps and epistatic interactions (10).

In practice, detecting genome-wide heterogeneity in recombination rates is challenging. Empirical approaches require building linkage maps through involved procedures such as sperm typing or multi-generational genetic crosses (11). While these are often the most powerful methods for detecting recombination, they are costly and time consuming. With the recent influx of large-scale sequencing data, alternative algorithmic approaches to inferring recombination rates from bulk genomic data have become a focus of attention. These methods, while often faster, have come with their own set of technical challenges. In order to detect patterns and rates of recombination along a genome, model-based algorithms infer properties of the ancestral recombination graph (ARG), an exercise which can be prohibitively computationally expensive on large datasets (12). This problem has driven the development of methods which use either a variety of summary statistics built on quantities such as the distribution of pairwise differences (13), or only compute partial or composite likelihoods, such as LDhat and its sister LDhelmet, two of the most widely-used model-based methods (14, 15). Even with this relaxation, these methods can take a matter of days to run on realistically sized sequence data.

In this paper, we present a method that takes advantage of novel summary statistics based on topological features of the genomes in a given sample to quickly and accurately provide estimates of recombination rate heterogeneity. Our method differs from existing model-based methods in that it is based solely on distances between sequences and consequently scales significantly better on large datasets. We find that a topological data analysis (TDA)-based approach greatly increases feasibility of the inference problem and implicitly ties genetic distances to modern models of population genetics via the coalescent (see *Coalescent Intuition for Topological Statistics*).

Recently, Camara et al. demonstrated the utility and efficiency of topological data analysis to inference of recombination rates (16), benchmarked against the methods of Hudson and Kaplan, Myers and Griffiths, and Chan et al. (15, 17, 18). They focused on a topological feature known as the first Betti number (*β*_1_, explained in *Background on Topological Data Analysis*), which captures the number of cycles in an ARG, the canonical graphical representation of recombination events. We have found that another topological feature of lower dimension, which we call *ψ*, is a better predictor of recombination rate in genomes. Moreover, *ψ* and *β*_1_ used in tandem provide much more accurate estimates than previous TDA-based methods. We investigate these two topological features and their relationships to evolutionary quantities of interest including recombination rate as well as coalescent tree length and describe a method of estimating recombination rates from genome samples using these features. We then compare the performance of our estimator on whole-genome data to LD-Helmet; we find that our results justify the use of the TDA estimator as a rapid approximation method.

## Background on Topological Data Analysis

Topological data analysis (TDA) is a new branch of statistics that applies tools from algebraic topology to describe the shape of data (19–23). TDA has been successfully applied to a range of applications in biology, including the study of breast cancer transcriptional data for the discovery of a cancer subgroup (24), the construction of phylogenetic trees for analyzing tumor evolutionary patterns (25), and the detection of intrinsic structure in neural activity (26). In this section, we briefly review the relevant TDA methodology that we apply to the study of recombination. (For a more detailed review of TDA applications in genomics, see (27).)

TDA associates topological summaries to data sets using a novel invariant called persistent homology (19–23). We begin by giving a brief explanation of homology; note that herein, homology refers to a mathematical concept rather than a biological one. Homology refers to a family of ways of associating an algebraic object (i.e., a vector space) to a geometric object. For example, the zeroth homology, *H*_0_, is associated with the number of unique connected components (0-dimensional holes) in the data, such as independent points or groups of connected points. The first homology, *H*_1_, captures cycles or loops (1-dimensional holes) in the data, the second homology, *H*_2_, captures voids or hollow spheres (2-dimensional holes) in the data, and so on. The rank of the *i*^*th*^ dimensional homology group is known as the *i*^*th*^ Betti number, denoted *β*_*i*_, and roughly speaking it encodes the number of *i*-dimensional holes in the dataset (19–23).

For example, a figure-8 consists of a single connected component (all the points in the boundary of the figure are connected) and two loops, so for this shape *β*_0_ = 1, *β*_1_ = 2, and *β*_*k*_ = 0 for *k* > 1. In contrast, a basketball is one connected component (all points on the surface are connected) with one hollow sphere and no loops so its associated Betti numbers are *β*_0_ = 1, *β*_1_ = 0, *β*_2_ = 1, and *β*_*k*_ = 0 for *k* > 2. To see why *β*_1_ = 0 for this example, consider any loop on the basketball and fix some point *p* on the surface of the ball. Without breaking the loop, it’s possible to slide the loop in a continuous manner (while remaining on the ball) towards *p* until eventually the loop contracts to *p*. Consequently all loops on the basketball are trivially equivalent to a point, i.e. *β*_1_ = 0.

While shapes and surfaces have well-defined homology groups, computing the homology of data is less straightforward. If one were to directly compute the homology of a dataset consisting of N distinct data points directly, the resulting Betti numbers would be *β*_0_ = *N* and *β*_*k*_ = 0 for *k* > 0, since the data has N connected components (that is, distinct points) and no higher dimensional holes. In this context the Betti numbers are clearly non-informative. Consequently, in data-driven tasks we instead seek to uncover the underlying shape on which the data lie and then compute the homology of that shape (19, 21, 23). Algorithmically, this procedure is carried out by assigning a combinatorial model of a space, called a *simplicial complex*, to the data.

Given a dataset *X* = {*x*_0_, *x*_1_, …, *x*_*N*_} living in some metric space (*S, d*_*S*_) and a distance parameter ∊ > 0, we think of simplicial complexes as an organized set of vertices (0-simplices), edges (1-simplices), triangles (2-simplices), tetrahedra (3-simplices), and higher dimensional simplices. The vertex set of the simplicial complex is the collection of data points {*x*_0_, *x*_1_, …, *x*_*N*_ }. Moreover, if 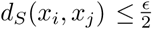, then the edge connecting *x*_*i*_ to *x*_*j*_ is included in the simplicial complex of X with respect to ∊. Similarly, if 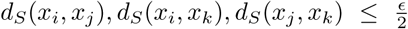, then the filled triangle with vertices *x*_*i*_, *x*_*j*_, *x*_*k*_ is contained in the complex (19, 21, 23). In general, a simplex is included in the simplicial complex representation of *X* with respect to ∊ if the vertices of the simplex have pairwise distance 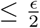. This simplicial complex is known as the Vietoris-Rips Complex, and it is the way in which we construct a simplicial complex from a set of genomes throughout this paper (where the Hamming distance provides the metric) (19, 21, 23). For a more rigorous definition of simplicial complexes, see *Supporting Information* and (21).

One can now compute the homology of the simplicial complex representation of a dataset to get non-trivial topological summaries. The main issue with this procedure is that it is sensitive to the choice of scale parameter ∊. The idea of persistent homology is to avoid making a choice of ∊ and instead track how the homology changes as ∊ varies (21, 28). For each dimension, persistent homology yields a collection of “homological features” which appear at some parameter value ∊_0_, known as the birth time, and disappear at some parameter value ∊_1_ ≥ ∊_0_, known as the death time. These features are often represented in a barcode diagram, where each bar represents an *i*-dimensional hole which persists within the interval [∊_0_, ∊_1_], as shown in Figure 1a (19, 28). One can think of persistent homology as an extension of hierarchical clustering for higher dimension homology groups, where dimension 0 persistent homology is analogous to single linkage clustering (19). See Figure 1b for an example of persistent homology applied to an arbitrary dataset via the Vietoris-Rips complex.

**Fig. 1.**
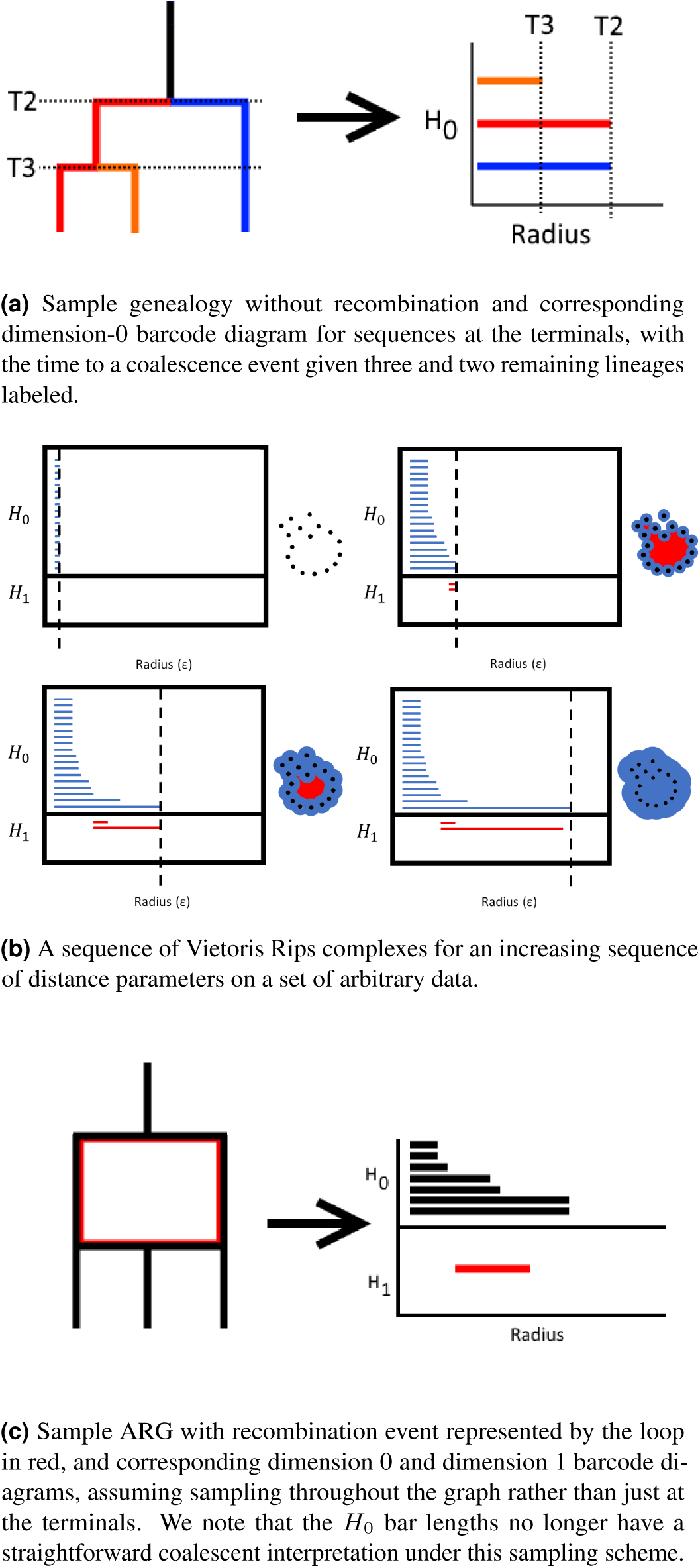
Persistent homology applied to genealogies in the absence and presence of recombination.

Intuitively, in the presence of recombination events the genealogy contains loops which correspond to dimension 1 homological features. This hypothesis was explored in certain cases in (16); the loops do not appear in the absence of recombination and consequently, *β*_1_ can be used to predict recombination rate. This is illustrated in figure 1c (although we note that in the context of standard coalescent assumptions, this connection is more complicated than it appears; see *Explaining* ***β***_**1**_ for details). One of the main discoveries we describe is that in fact the mean barcode length in dimension 0, denoted *ψ*, is an even more accurate predictor of recombination rate than *β*_1_. We note that each bar in the dimension 0 barcode diagram corresponds to an individual in the sample population, and the bar lengths correlate with distance between the individual, or cluster of individuals, and its closest neighbor in Hamming space. Unexpectedly, this measure of distance between individuals is positively correlated with recombination. An explanation of this behavior is located in *Coalescent Intuition for Topological Statistics*.

## Methodological Overview

We sought to identify topological summary statistics (using *β*_1_ as a baseline for comparison) that can serve as features for algorithms to perform recombination rate inference. Utilizing simulated data, we computed a variety of topological summaries of dimensions 0, 1, and 2 from the Hamming distance matrix between sequences.

The results of our LASSO regression indicated that the topological features with the highest predictive power for recombination rate are, in order: 1) the average dimension 0 barcode length (*ψ*), 2) the first Betti number (*β*_1_), and 3) the variance of the dimension 0 barcode lengths (Φ). We then used a nonlinear combination of these three topological statistics to build a novel TDA-based model for recombination rate inference, the Topological REcombination Estimator (TREE).

We used simulated data to perform an initial validation of the model. For a more serious validation, we applied TREE to 22 full genome assemblies from the RG Drosophila population (see *Methods* for more details) and compared its performance to *ρ*_*ph*_, the recombination rate estimator introduced by Camera et. al in (16), and to LDhelmet. We also benchmarked TREE on a much larger dataset of Arabidopsis genomes, consisting of 1,135 individuals and up to 50k SNPs.

### Coalescent Intuition for Topological Statistics

The main topological statistics of interest here are *β*_1_, the first Betti number, and *ψ*, which corresponds to the mean bar length in the dimension 0 barcode diagram. In order to relate these to the biological process of recombination, we will use the language of coalescent theory. For a detailed introduction to the field, see (29). We note that our approach differs from the recent considerations of Lesnick, Rabadán, and Rosenbloom (30) in that we consider a coalescent model with branch lengths and model *H*_0_ behavior. Furthermore, we assume a more restrictive sampling regime where only sequences at contemporaneous terminals of the graph are known, as opposed to sequences all along the genealogy.

#### Explaining ψ

We will provide a heuristic argument that the value of *ψ* is elevated in the presence of recombination by demonstrating the desired behavior at the recombination rate extremes. First, we claim that in the absence of recombination, the distribution of *H*_0_ feature lengths corresponds to the mutation scaled distribution of branch lengths in the coalescent tree of the sample, as shown in figure 1a. Since there is a single, fixed genealogy which describes all positions within the sequence, it is sufficient to calculate the expected length of the coalescent tree and divide by the sample size. Assuming an idealized diploid population of size *N*, large, and that we are sampling *K* individuals with *K* sufficiently small relative to *N* that multiple merger events are rare, the expected waiting time between coalescence events is 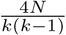 generations (31, 32), where *k* is the number of remaining lineages. The full coalescent tree is then made of each *k*^*th*^ interval *k* times, for the number of remaining lineages at that time. Summing over all these segments and dividing by the sample size gives us the following:

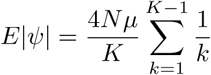

where *μ* is the per-generation mutation rate. Notably,this is equivalent to the expected number of segregating sites divided by the sample size per Watterson’s estimator (31).

We now show that in the infinite recombination limit, the expectation for *ψ* is strictly larger than in the case where there is a single fixed genealogy. If there is free recombination, every site in the sample has an independent genealogy which we will average over, so all the bars must be of the same length (Figure 2). In other words, the expected value of *ψ* becomes the scaled average coalescence time for two randomly sampled individuals in the current generation. This is simply 2*μN*. All that remains is to show that 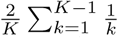 is less than 1. Since the partial sum is bounded by log(*K* – 1) + 1, this holds for all values of *K* larger than 5. This additionally suggests that the variance in the length of the *H*_0_ barcodes features should decrease as the recombination rate increases, which we observe in simulations. By integrating information both about the length of the coalescent tree as well as the distribution of pairwise differences averaged over multiple topologies, *ψ* can be viewed as capturing distortions in the expected amount of independent evolution between samples that occurs when sequences contain multiple discordant gene genealogies.

**Fig. 2.**
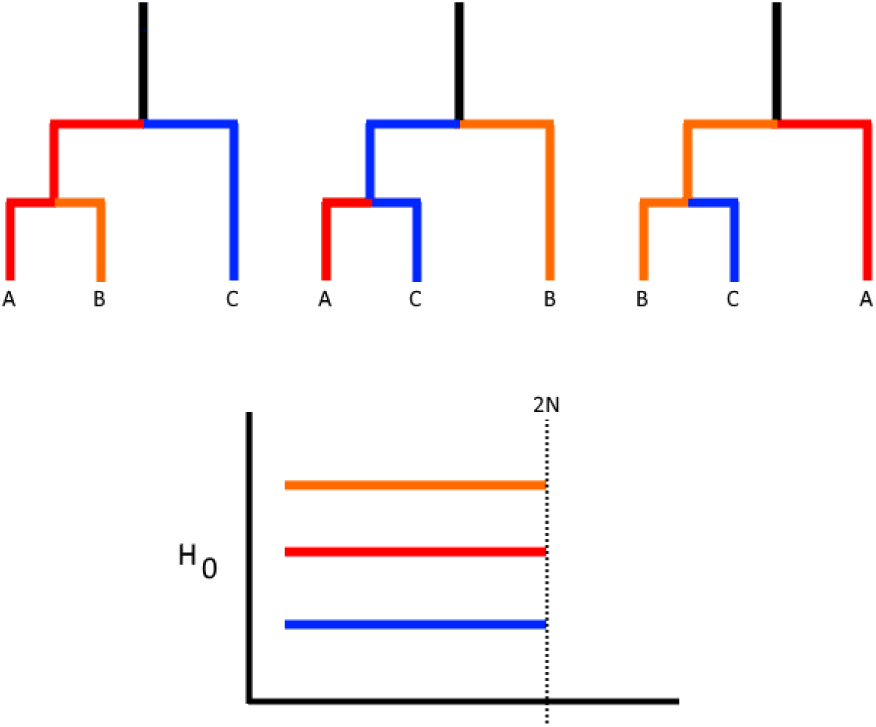
By averaging over multiple genealogies, the barcode of *H*_0_ features approaches identical bars of length equal to the expected pairwise coalescence time.

#### Explaining β_1_

We suggest in addition that the standard intuition for the use of *β*_1_ to detect cycles in the ARG (presented in Camera et al. and Chan et al. (16, 33), as well as here in *Background on Topological Data Analysis*), potentially over-simplifies the rather complicated relationship between recombination events and features in the *H*_1_ barcode. For this, we will consider Čech complexes, rather than Vietoris-Rips complexes, as they are closely related (22), and the Čech complex construction allows holes to be formed given only three points (see Figure 3a). Given the graph in Figure 3b, it is clear that if one were to sample the sequences at every node, there would be an *H*_1_ feature observed which corresponds precisely to the hole in the graph. However, in many genetic studies, samples of the common ancestors of present-day sequences do not exist. If we restrict our data to the sequences at the terminals, cycle detection with *β*_1_ becomes a function of mutation heterogeneity along the graph, and is in fact impossible if we are given only the true coalescent distances between samples along the genealogy. To see this, take terminals *a* and *c* from the graph. By hypothesis, the amount of time between them and their most recent common ancestor (MRCA) is the same, which we will call *L*. It follows that the minimum radius such that balls around these points would intersect is *L*. Then, for the triple intersection of balls around the terminals to be empty, *b* must be a distance greater than *L* from the MRCA. However, each portion of its genome has certainly experienced the same amount of time since the MRCA regardless of recombination history. Therefore, we require that there be a more than expected amount of mutations generated along the path to *b* in order for this event to be detected, given the actual sequence data. It still holds true that *β*_1_ reflects only these recombination events, assuming no sequence convergence and infinite sites, but this sampling reality may bias *β*_1_ detection in subtle ways. This also implies that the length of the *H*_1_ bar will not necessarily be indicative of features of the actual cycle in the graph, as it will increase in length as additional mutations are placed on the lineage leading to *b*, even if the cycle itself is untouched. We find via simulations (see supplement *Filtering* ***β*_1_**) that filtering small *H*_1_ bars only hurts our inference capabilities, as we would then expect.

**Fig. 3.**
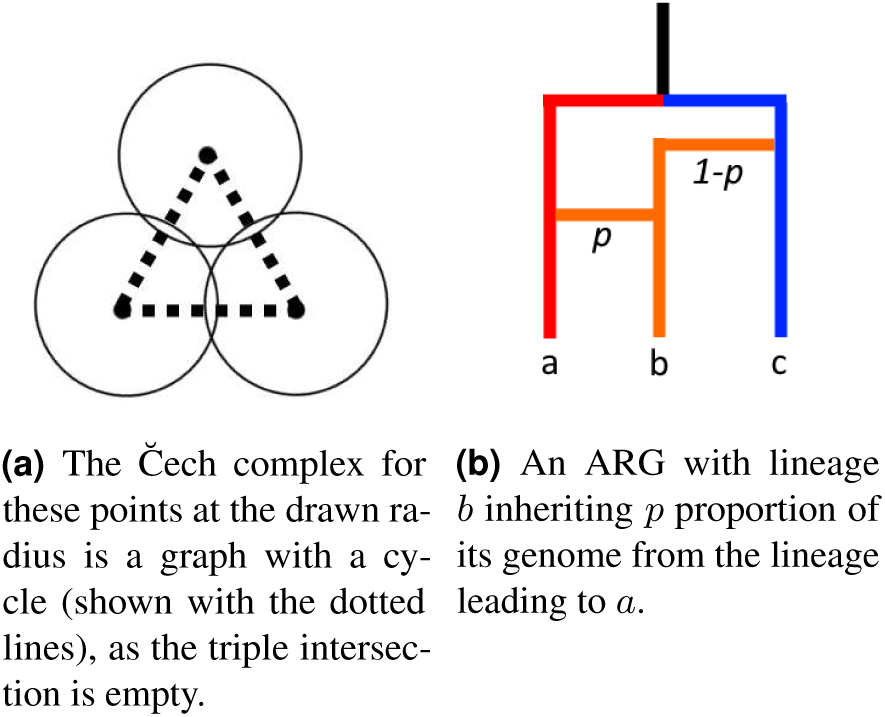
Given the true coalescent distances between the terminals *a, b, c*, a recombination cycle will not be detected in the manner shown in panel 3a, and so will not generate an *H*_1_ feature in the čech complex at any radius.

#### Combining ψ and β_1_

These explanations for the behavior of *ψ* and *β*_1_ also implies differences in behavior between *ψ* and *β*_1_ under different population models. For example, while rapid demographic changes such as exponential population growth will distort *ψ*-based estimation (Figure 4), *β*_1_ counts features generated by recombination at a rate independent of the underlying tree structure, since the relative distances to the MRCA are unchanged with multifurcations. On the other hand, we find empirically *ψ* is very robust (especially compared to *β*_1_) to perturbations in the form of missing data, which serve only to minorly rescale the *H*_0_ bars on average. These differences suggest that a reliable predictor should incorporate both features. A more formal follow-up to the behavior of these statistics under different models of demography and selection, as well as further characterization of the behavior of *ψ* using the sequentially Markov coalescent (SMC) model (34) will be conducted in future work.

**Fig. 4.**
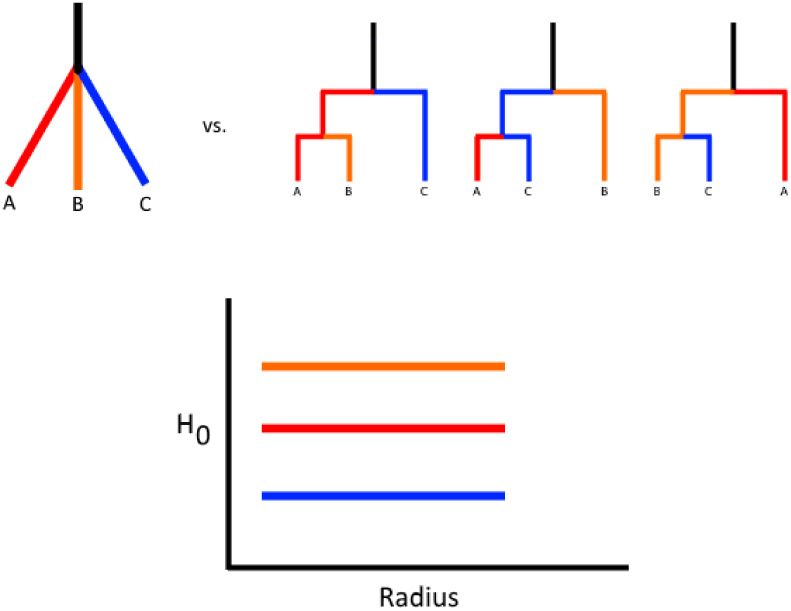
Exponential population expansion creates multiplemerger events and shrinks internal branches. This can give a similar signal in *H*_0_ as increased recombination, but does not change cycle detectability via *β*_1_ in the ARG.

## Results

### Coalescent Simulations

We generated simulated data for an idealized population of fixed size *N*_*e*_, with per-generation crossover rate *r* and mutation rate *μ*. Given this, we ran regression analyses to discover relationships between various topological summary statistics and *ρ* = 4*N*_*e*_*r*, the population recombination rate. The topological statistics analyzed were the mean and median bar lengths, variance of bar lengths, total number of bars, and number of bars above varying noise-filtering thresholds for dimensions 0, 1, and 2.

Our intention was to test various topological features as predictors of known recombination rates and to demonstrate comparable performance of these features to a comparator method, LDhelmet. However, given the computational bottlenecks inherent to LDhelmet, we were unable to run this software over the full set of >3600 alignments. We are nevertheless satisfied in that the parameters to be estimated are known from simulation, and so we save the use of LDhelmet for our empirical analyses where the truth is unknown.

As a preliminary analysis, the weight vectors of LASSO regression models provided insight into which barcode statistics were the strongest predictors of *ρ*. The results of this analysis showed that two key topological features, *β*_1_ and the mean dimension 0 bar length, which we will denote as *ψ*, correlate strongly with *ρ* given a constant sample size *n* > 10. Thus, these topological summaries became the main foci of our work. Moreover, we found no correlation between our barcode statistics and *θ* = 4*N*_*e*_*μ*, the population mutation rate, as expected, since changes in *θ* with a constant *N*_*e*_ only linearly rescale the distances.

In studying the information content of these various statistics, we found *ψ* stood out as an even better predictor of recombination rate than *β*_1_, and performed better on its own in predicting recombination rates from simulated datasets. Figure 5 demonstrates the relationships between *ρ* and *β*_1_, and between *ρ* and *ψ* for sample sizes *n* = 50 and *n* = 140. We note that both topological summaries exhibit an exponential relationship with recombination rate, and that the relationship is tighter for *ψ* than *β*_1_, especially for smaller sample size.

**Fig. 5.**
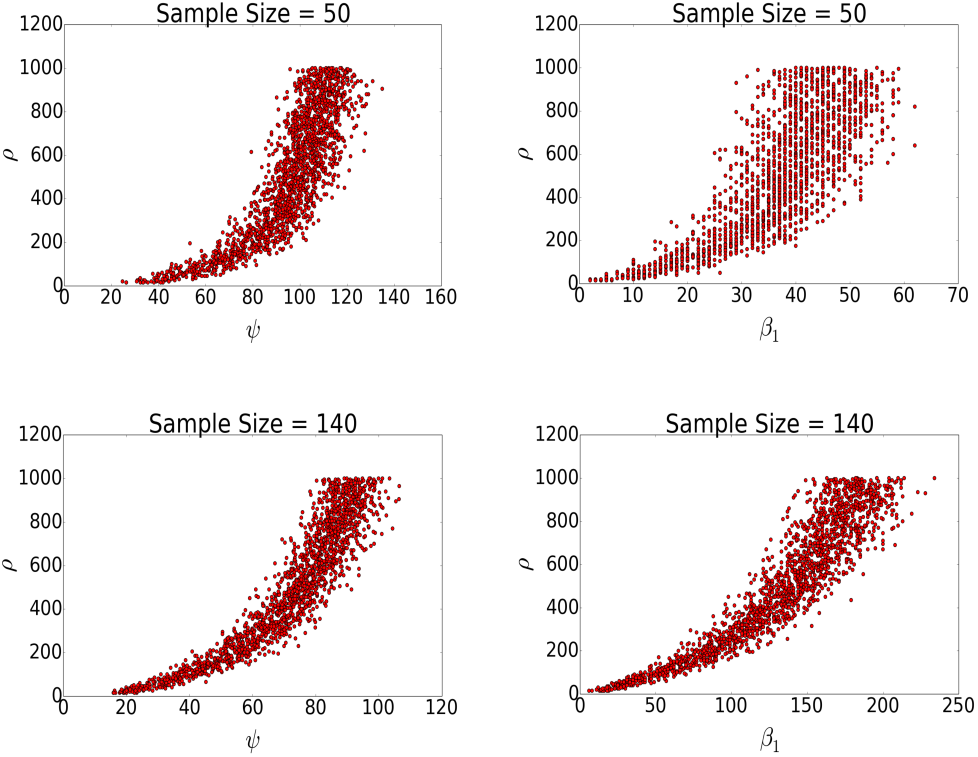
The relationship between *ψ* and recombination rate (left), and *β*_1_ and recombination rate (right) for a simulated dataset with fixed sample size *n* = 50 (top) and *n* = 140 (bottom).

As a baseline, we fit our simulated data with a fixed sample size of 160 to the *β*_1_ based model of Camara et al. (2017), *ρ*_*ph*_ (the “ph” stands for persistent homology). This model is given by the equation

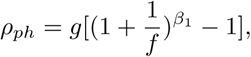

where the parameters g and h are coefficients related to sample size and are independently calculated for a given dataset. The best fit over the simulated range of recombination rates simulated is *R*^2^ = 0.86. We found a relationship between *ρ* and *ψ* with *R*^2^ = 0.90, suggesting that this new parameter has comparable or greater power for predicting population recombination rates.

We demonstrated that the two parameters *ψ* and *β*_1_ are differentially stable under violations of assumptions about the data. Notably, *ψ* is robust to large amounts of missing data, whereas *β*_1_ is robust to rapid changes in population size. The former we show empirically: *ψ* maintains its relationship with *ρ* with *R*^2^ = 0.76 with 10% of each sequence missing as a tandem indel, while *β*_1_ loses this relationship quickly with *R*^2^ = 0.036 under the same missing data scheme. The assumption of 10% missing data is somewhat conservative; nextgeneration sequencing data sets have less missing data, but this figure is realistic for many older sequencing data sets. As one would expect, we note that performance of each method declines as missing data become more prevalent (See supplement *Missing Data*). However, if missing data is located randomly throughout the genome, it is unlikely to bias relative measures of recombination rates within a dataset. While *β*_1_ is expected to be robust to rapid changes in population size since this does not change the number of cycles in the ARG, *ψ* will be more sensitive to these changes as rapid demographic expansion generates multifurcations in the genealogy which converge to the same branch lengths as in the infinite recombination case.

### TREE Model

We implemented several machine learning and regression models to build an accurate and robust model relating topological summaries to *ρ*, including LASSO, polynomial regression, exponential regression, and linear regression. We varied model parameters and input features (subsets of the aforementioned barcode statistics) across each learning algorithm.

**Fig. 6.**
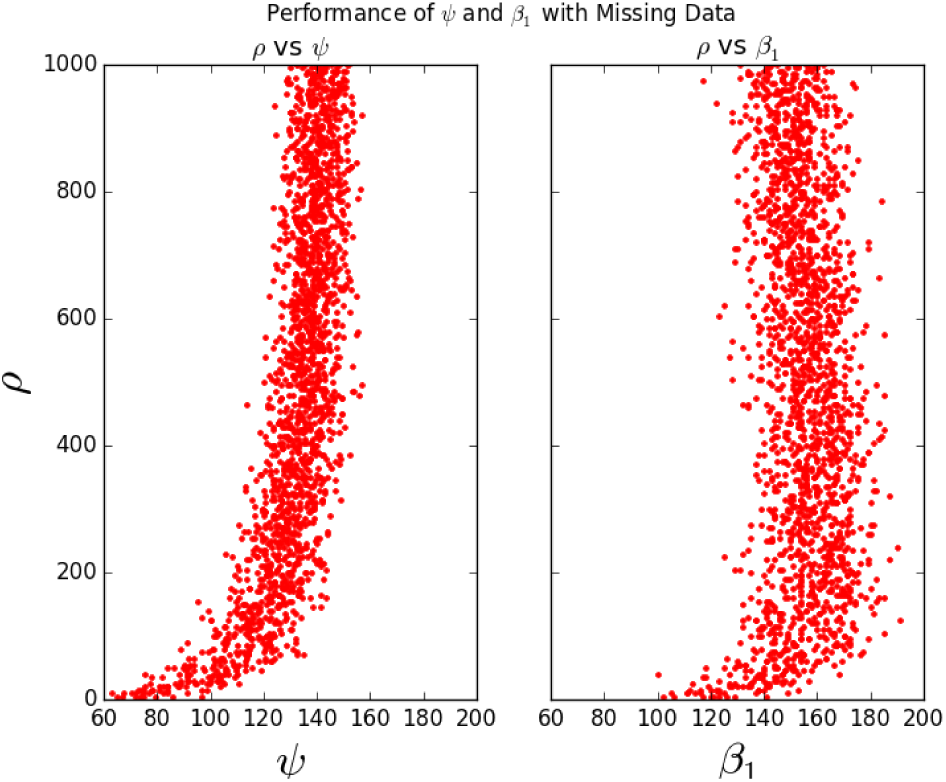
*ψ* continues to be accurate under a missing data scenario (*R*^2^ = 0.76 with 10% of the data missing in large blocks) while the accuracy of *β*_1_ under the same scenario drops to *R*^2^ = 0.036.

A subset of model comparison results are presented in Table 1, where we show the *R*^2^, or goodness-of-fit measure, values for the model 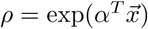, where 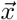 corresponds to a vector of different barcode statistics. We ran the model separately on datasets generated with varying sample sizes, as well as on a dataset consisting of simulations of varying population size. An *R*^2^ value close to 1 signifies a nearly perfect model, whereas *R*^2^ near 0 indicate that the model has low predictive power.

**Table 1.** *R*^2^ values for the exponential regression model for different feature inputs and sample sizes.

| Sample Size | $\psi$ | $\beta_1$ | $(\psi^2, \beta_1^2, \psi, \beta_1, \Phi)$ |
| --- | --- | --- | --- |
| 25 | 0.724 | 0.456 | 0.792 |
| 50 | 0.847 | 0.749 | 0.894 |
| 75 | 0.883 | 0.827 | 0.927 |
| 95 | 0.898 | 0.862 | 0.941 |
| 140 | 0.909 | 0.887 | 0.959 |
| mixed | 0.414 | 0.164 | 0.851 |

The results show that *ψ* is an overall stronger predictor than *β*_1_, and as expected, recombination rate is predicted more accurately for higher sample sizes. Importantly, *β*_1_ fails as a predictor in the case of small sample size, while *ψ* maintains decent predictive power for sample size as low as 25.

A thorough comparison of the different model outputs showed that an exponential model in *ψ*^2^, *β*_2_, *ψ, β*_1_, and the variance of the dimension 0 bar lengths, which we will denote Φ, is the best predictor for *ρ*. While we also tested more topological features as inputs to different models, the increase in *R*^2^ values was negligible in comparison to the increased risk of over-fitting.

Summarizing, we propose the following Topological REcombination Estimator (TREE) model:

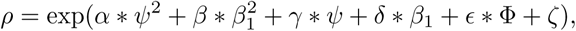

where

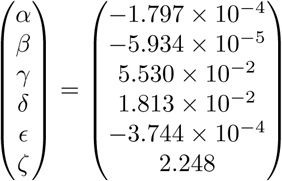

Figure 7 shows TREE’s performance on blind testings set for sample sizes of 50 and 140. While the model preforms best when restricted to datasets corresponding to high sample size, TREE was trained on mixed sample size data in an attempt to make the model as robust and have as little bias as possible.

**Fig. 7.**
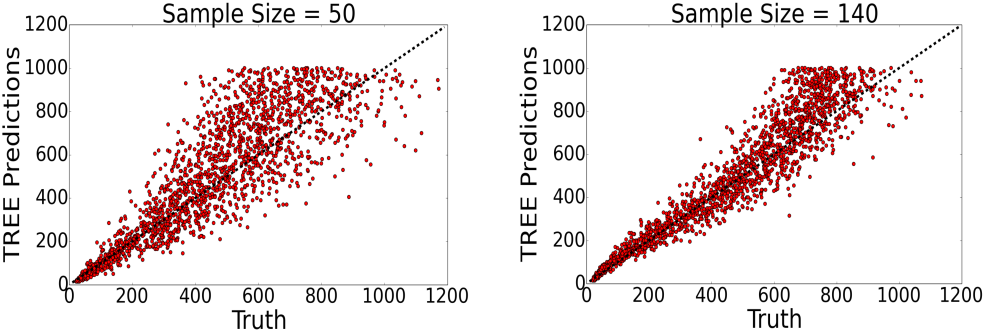
TREE predictions on testing sets compared to the true recombination rate for sample size = 50 (left) and sample size = 140 (right). The dotted line corresponds to perfect predictions.

To analyze the robustness of the model we preformed fivefold cross validation. The results are presented in table 2, and they show that the model maintains high accuracy regardless of the training set.

**Table 2.** Five-fold cross validation *R*^2^ values for the full model.

| Sample Size | $k = 1$ | $k = 2$ | $k = 3$ | $k = 4$ | $k = 5$ |
| --- | --- | --- | --- | --- | --- |
| 25 | 0.797 | 0.769 | 0.738 | 0.696 | 0.786 |
| 50 | 0.857 | 0.901 | 0.905 | 0.884 | 0.881 |
| 75 | 0.901 | 0.934 | 0.906 | 0.925 | 0.928 |
| 95 | 0.938 | 0.934 | 0.944 | 0.936 | 0.952 |
| 140 | 0.956 | 0.963 | 0.949 | 0.958 | 0.961 |
| mixed | 0.849 | 0.847 | 0.849 | 0.852 | 0.858 |

### Empirical Analysis

We compared the performance of TREE on empirical datasets to a widely used estimator, LDhelmet v1.9. We first benchmarked each method on a large genomic dataset from the Arabidopsis 1000 Genomes project consisting of 1,135 samples. We used subsets of 1k, 10k, and 50k SNPs in 100 SNP windows from the total dataset in order to test the computational speed of each program. We found that LD-helmet cannot process datasets with greater than 50 samples, failing to complete the first step of the likelihood table computations for the smallest of these datasets. However, TREE was able to process each dataset in full within a reasonable time frame (Table 3). By sub-sampling these data, we can illustrate one advantage of TREE’s ability to analyze this quantity of individuals. We find that as the number of samples is increased from 20%to 50% to 100% of the full set, the distribution of recombination events along the genome shifts such that the top 10% of windows contain 17.4%, 19.8%, and 23.9% of events respectively, as the signal of hotspots grows more pronounced. In addition, the importance of efficiency is underscored by the fact that as more samples are included, more SNPs are realized in the data, and more windows are required for a full analysis. As we could not compare recombination estimates on the full Arabidopsis genome due to LDhelmet’s processing times, we turned to a smaller Drosophila dataset with 22 samples and >22Mbp. For these datasets, LDhelmet takes on the order of hours to complete a run over a single chromosome, whereas TREE terminates on the order of minutes and in all cases finished running in under one hour.

**Fig. 8.**
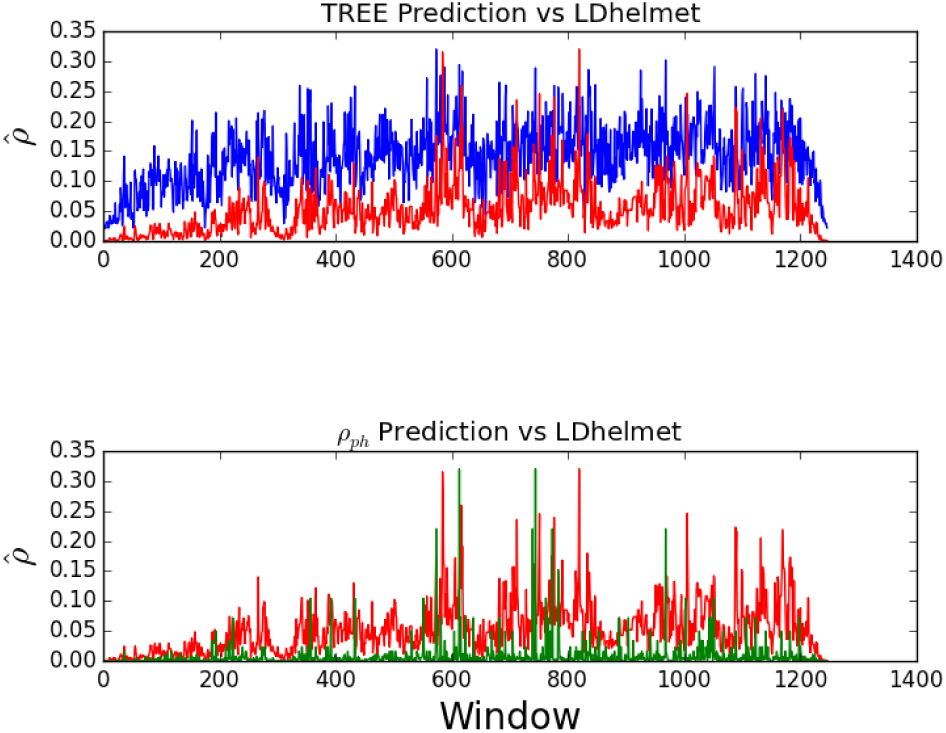
Relative accuracy of TREE and Camara’s *ρ*_*ph*_ with respect to LDhelmet. The blue line plot represents TREE, the red represents LDhelmet, and the green in the second panel represents *ρ*_*ph*_. Due to issues with *ρ*_*ph*_ orders of magnitude more recombination rate variation than LDHelmet, we rescale the estimate given by *ρ*_*ph*_ to the exact range of LDHelmet. For TREE, we only multiply by a uniform window length conversion factor of 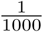.

**Table 3.** Benchmarking TREE’s runtime on a large dataset (1135 Arabidopsis individuals)

| Number of SNPs | Runtime (hours) |
| --- | --- |
| 1k | 0.521 |
| 10k | 5.556 |
| 50k | 27.866 |

We first take a broad look at the relative performance of TREE to LDhelmet on genomic datasets, looking for concordance in predicting an increase, decrease, or no change between each window of our analysis. We find that TREE is concordant with LDhelmet 69.2% of cases where *ρ* increases and in 69% of cases where *ρ* decreases. We note that since LDhelmet applies a smoothing with TREE does not, we cannot directly compare the accuracy with which TREE generates adjacent windows of identical recombination rate.

To characterize TREE’s behavior in these cases, we looked at the magnitude of the difference between the TREE prediction and LDhelmet’s. We found that in the cases where LDhelmet predicts no change in *ρ*, TREE’s predicted change is less than 0.05 72.5% of the time, less than 0.01 18.1% of the time, and less than 0.001 1.8% of the time. These results suggest that TREE is good at detecting large changes in recombination rate, and is thus well-suited for hotspot detection. However, it can have difficulty differentiating between subtler changes in recombination rate ranging in magnitude between 0 and 0.01.

Finally, we looked directly at the correlation between the absolute predicted values of TREE and LDhelmet for the most fine-grained comparison. We used two measures of correlation the Kendall-Tau rank test, and Spearman’s *ρ*. With the exception of chromosome arm 2L, Kendall’s Tau between TREE estimates and LDhelmet estimates ranges between 0.067 and 0.241 with P-values less than 0.0001. Similarly, Spearman’s *ρ* ranges between 0.097 and 0.346 with P-values less than 0.0002. These positive correlation coefficients and low p-values suggests global agreement between LDhelmet’s and TREE’s rankings of recombination rates across sliding windows, indicating that TREE is useful in detecting global hotspots of recombination.

Chromosome X and arm 2L are outliers here, as their correlation coefficients are substantially lower and associated Pvalues substantially higher than the remaining chromosome arms in the dataset (Table 4). Despite attempts to discover why these two chromosomes in particular are outside of the average ranges of performance, we were unable to find a compelling reason. It may be that each of these chromosomes have many more cases of subtle recombination rate changes between 0-0.1, such that LDhelmet’s estimates are most different from TREE’s.

**Table 4.**
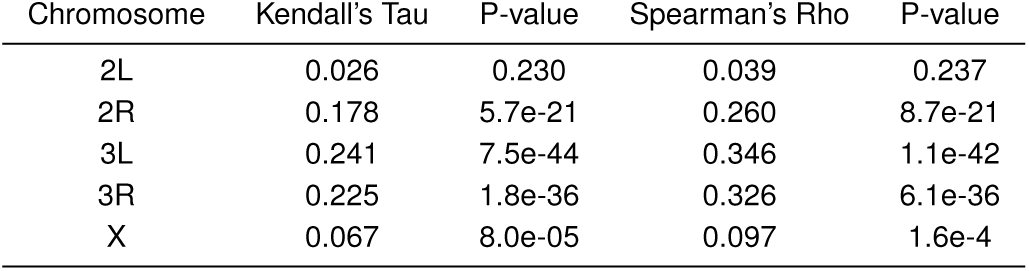
Comparison of TREE to LDhelmet’s *ρ* estimates

| Chromosome | Kendall's Tau | P-value | Spearman's Rho | P-value |
| --- | --- | --- | --- | --- |
| 2L | 0.026 | 0.230 | 0.039 | 0.237 |
| 2R | 0.178 | 5.7e-21 | 0.260 | 8.7e-21 |
| 3L | 0.241 | 7.5e-44 | 0.346 | 1.1e-42 |
| 3R | 0.225 | 1.8e-36 | 0.326 | 6.1e-36 |
| X | 0.067 | 8.0e-05 | 0.097 | 1.6e-4 |

We also compared the model of Camara et al. to LDhelmet in the same framework to discover its relative performance, in order to discover whether the addition of the feature *ψ* is necessary for greater accuracy. We found that the *ρ*_*ph*_ model in *β*_1_ alone is dramatically less concordant with LDhelmet estimates across all windows than is TREE (Table 5). For each chromosome arm analyzed using *ρ*_*ph*_, the rank coefficients of Kendall’s Tau or Spearman’s *ρ* are less than 0.01, sometimes negative, and not statistically significant. This indicates poor concordance between the two methods and, in some cases, disagreement, suggesting that the addition of *ψ* to an estimator of recombination rate substantially improves accuracy as well as computation time compared to LDhelmet. We see evidence of the differences in prediction between TREE and LDhelmet as well as *ρ*_*ph*_ in Figure 7, which shows that TREE approximates the estimates of LDhelmet across the entire span of the chromosome without rescaling, whereas *ρ*_*ph*_ fails to capture similar detail to TREE and requires an informed re-centering to the range of LDHelmet to be competitive.

**Table 5.** Comparison of *ρ*_*ph*_ to LDhelmet’s *ρ* estimates

| Chromosome | Kendall's Tau | P-value | Spearman's Rho | P-value |
| --- | --- | --- | --- | --- |
| 2L | 0.017 | 0.446 | 0.02 | 0.430 |
| 2R | -0.104 | 4.1e-8 | -0.154 | 5.0e-8 |
| 3L | 0.005 | 0.754 | 0.008 | 0.745 |
| 3R | -0.002 | 0.879 | -0.003 | 0.918 |
| X | 0.013 | 0.449 | 0.019 | 0.435 |

**Table 6.**
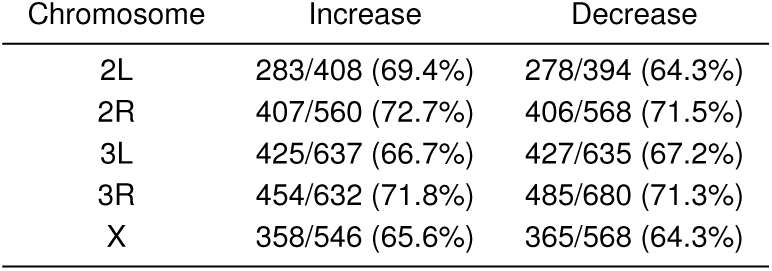
Compared of TREE to LDhelmet by chromosome arm in terms of the change in *ρ*

## Discussion

We have discovered a new feature of the distribution of genomes in Hamming space, which we denote *ψ*, that improves the performance of topological estimators of recombination. While the field of TDA is in its infancy, our work provides a novel demonstration of the power of persistent homology-based estimators for fundamental questions in evolutionary biology. Notably, our feature is related to biologically-meaningful quantities in coalescent models; this is some of the first work we are aware of to make such a tight connection between TDA estimators and coalescent theory.

Our *ψ*-based approach to recombination rate inference is able to quickly scan large genomic datasets for regions of recombination rate heterogeneity. Due to its speed, it can serve as a first-pass estimate of recombination rate variation prior to targeted use of much more computationally expensive inference methods. While *ψ* itself can potentially be influenced by distortions to the genealogical structure of a sample, it is naturally complemented by higher dimensional topological features (namely *β*_1_) of the data explored in prior work (16, 33), while maintaining accuracy in the face of missing data which confounds *β*_1_-only methods.

Similarly to how *ψ* can supplement and guide usage of evolutionary-model driven methods, *ψ* can also add a degree of finer-scale detection and biological intuition to topology-driven methods, bringing us closer to bridging the gap between population genetics and persistent homology. The differing behaviors of *H*_0_ and *H*_1_ derived statistics on genomic data also point towards the potential of TDA as a source for summary statistics which can tease apart the signatures of demography, selection, or population structure, a fundamental goal of population genetics. The behavior of *ψ* also suggests that topological quantities could be merged with a fully coalescent model of recombination, as a more rigorous SMC-based modeling could make explicit predictions for the distribution of the effect sizes of a single recombination events on *ψ*. In addition, the ordering of the *H*_0_ features by duration of persistence hints at additional connections between barcodes and the look-down construction of the coalescent (35, 36). We will explore these avenues in a future theoretical treatment of these statistics.

**Table 7.** Comparison of *ρ*_*ph*_ to LDhelmet by chromosome arm in terms of the change in *ρ*

| Chromosome | Increase | Decrease | No Change |
| --- | --- | --- | --- |
| 2L | 199/408 (48.77%) | 215/394 (54.57%) | 8/116 (6.90%) |
| 2R | 133/560 (23.8%) | 128/568 (22.5%) | 64/118 (54.2%) |
| 3L | 290/637 (45.5%) | 283/635 (44.6%) | 12/204 (5.8%) |
| 3R | 298/632 (47.2%) | 321/680 (47.2%) | 3/78 (3.8%) |
| X | 243/546 (44.5%) | 262/568 (46.1%) | 30/411 (7.3%) |

Summarizing, we have shown that a combination of TDA and machine learning techniques can detect recombination rate heterogeneity in biological data faster than previously possible and with greater accuracy than previous TDA-based approaches. We demonstrate that while the behavior of 0dimensional barcodes has been previously ignored with respect to genealogical inference problems, these features are robust and increase the overall accuracy of inference compared to using 1-dimensional barcodes alone. Our coalescent analyses also suggest a promising future endeavor: building a fully coalescent-motivated model explaining the behavior of Betti numbers on distributions of genome sequences.

## Methods

### Topological Data Analysis

We created a pipeline which takes as input a sequence file in FASTA format, computes a Hamming distance matrix *D*_*H*_, uses *D*_*H*_ to extract the corresponding dimension 0 and dimension 1 persistent homology barcodes, and then calculates barcode summary statistics. The barcodes were computed using Ripser, a publicly available C++ package for computing Vietoris–Rips persistence barcodes, and all other computations were done in Python 2.7 (37). The scripts are available on our Github page at: https://github.com/MelissaMcguirl/TREE.

### Simulations

We simulated over 50,000 datasets of genetic sequence data with known recombination rates, each of length 1000 bp, using the programs ms and seq-gen (38). Given an idealized population, we varied the parameters for the population recombination rate *ρ* = 4*N*_*e*_*r*, sample size *n*, and population mutation rate *θ* = 4*N*_*e*_*μ* to capture a variety of mutation-recombination regimes in the data. The datasets had recombination rates varying from 0 to 1000 (that is, no recombination up to free recombination between all sites under the population recombination model *ρ* = 1000).

For each population we computed dimension-0 and dimension-1 persistent homology barcodes using Ripser (37). We ran regression analyses to discover topological predictors for recombination and then confirmed that our method predicts *ρ* directly and does not predict a covariate such as *θ*.

### Model Selection

Initially, we applied polynomial regression of degree two, linear regression, LASSO regression, polynomial LASSO regression, and exponential regression using several different combinations of barcode statistics as inputs for predicting recombination rate using Scikit-learn (39). We sought the simplest model with the least number of parameters that was able to predict recombination rate while still maintaining high accuracy.

Each model was trained on a randomly selected subset of the input data, whose size was chosen to be 30% of the total dataset. The model was then tested on the remaining 70% of the input data, where *R*^2^ values were computed to access the goodness-of-fit of the resulting model. This process was repeated several times to test the robustness of the learned parameters with respect to different training sets.

Based on the *R*^2^ values, we were able to narrow down to three barcode statistics, *ψ* (average dimension 0 bar length), *β*_1_ (first Betti number), and Φ (variance of dimension 0 bar lengths), as inputs for an exponential regression model of the form

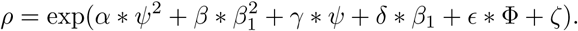

The coefficients (*α, β, γ, δ,* ∊, *ζ*) were determined via ordinary least squares from simulated training data in scikit-learn Python 3.0).

This model was tested on input data consisting of 53,461 barcode statistic files corresponding to simulations of varying sample size, mutation rate, and recombination rate. Five-fold cross validation was preformed for different sample sizes, and for the complete dataset.

### Empirical Analysis

We obtained 22 full genome assemblies from the RG population (from the African survey of *Drosophila melanogaster*) for our analyses. We ran LDhelmet and TDA in parallel and compared the mean estimates of recombination rate *ρ* in sliding windows of 500 SNPs. Since LDhelmet provides estimates of *ρ* between any two SNPs, whereas our method computes *ρ* within windows of 500 SNPs, we take the mean of every 500 *ρ* estimates from LDhelmet to compare to our windows. Moreover, since LDhelmet estimates *ρ* = 2×*N*_*e*_×*r* per base pair, and TREE predicts *ρ* = 4×*N*_*e*_×*r*, where *N*_*e*_ is the population size and r is the probability of a crossover event from ms, then we apply a uniform normalization of 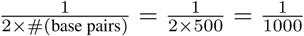 to the TREE predictions for comparison to LDHelmet.

We also ran Camara et al’s model using only *β*_1_ as a predictor within our sliding window framework to compare this to our method and LDhelmet. We looked at the results in 3 different ways: 1) in terms of absolute estimates compared to each other, 2) in terms of concordance in the change in *ρ* across windows (i.e. do both methods predict an increase or decrease in *ρ* in the same window), and 3) in terms of concordance of estimates above the 75th and 90th percentiles of the distribution of estimates. All analyses were done in Python 3.0, and scripts are available on our Github page: https://github.com/MelissaMcguirl/TREE.

We additionally included an analysis of 1,135 publicly available Arabidopsis genomes. We converted the raw VCF file to FASTA using a combination of bcftools and VCF-kit’s phylo fasta function, subsampling up to 50k SNPs in order to run the software within Stampede2’s 48 hour time limit. We ultimately used this dataset to benchmark the computational efficiency of TREE over LDhelmet, as LDhelmet is unable to process the full dataset with so many samples for us to use this as a comparator of accuracy.

## Acknowledgements

M.M. would like to thank John Wakeley as well as members of the Desai lab for helpful comments and discussion. The authors would like to thank Raul Rabadan and Juan Patino Galindo for very helpful feedback on the manuscript. D.P.H, M.R.M., and M.M. are supported by the National Science Foundation Graduate Research Fellowship Program under Grant Numbers DGE1610403, 1644760, and DGE1745303, respectively. A.J.B. was supported in part by NIH grants 5U54CA193313 and GG010211-R01-HIV and AFOSR grant FA9550-15-1-0302. Any opinions, findings, and conclusions or recommendations expressed in this material are those of the authors and do not necessarily reflect the views of the National Science Foundation.

## Supplemental Information

### Topological Data Analysis

Topology is a branch of mathematics that concerns itself with classifying spaces or objects that have the same ‘shape.’ Spaces are considered to be topologically equivalent if you can deform one into the other without breaking, tearing, or gluing. A topological invariant of interest in algebraic topology is the homology. Homology can be thought of as a family of ways to associate a vector space to a geometric object. For the scope of this paper it suffices to restrict ourself to homology with ℤ*/*2ℤ coefficients in dimensions 0,1, and 2. For simplicity we can think of the dimension 0 homology group as a representative of the connected components of a topological space, the dimension 1 homology group as a representative of the loops within a topological space, and the dimension 2 homology group as a representative of the void within a topological space. That is, the 0-th dimension homology group of an object is a vector space whose dimension is the number of connected components of that object, and similarly for higher dimensions.

Topological data analysis (TDA) lies at the intersection of algebraic topology, statistics, and data science. The main goal of TDA is to extract descriptive topological features from large, high-dimensional data sets and one of the primary tools for doing so is called, *persistent homology*.

Let *X* be a data set consisting of N data points living in some metric space (*S, d*_*S*_). Observe, *X* itself is simply a discrete set of points and thus it has no interesting homological properties beyond dimension 0. However, if each data point *x ∊ X* is replaced by a ball *B*_*r*_(*x*) = {*y* : *d*_*S*_(*x, y*) ≤ *r*} of radius *r* > 0 centered at *x*, then the union of these balls over all points *x* ∊ *X* yields a new topological space with non-trivial homology. Repeating this for a sequence of *r* values yields a sequence of topological spaces for which the homology can be computed. Analyzing how the homology changes across this sequence of topological spaces is the main idea behind persistent homology.

Formally, let *X* be a data set living in a metric space and choose *r* > 0. In order to compute the homology of X with respect to *r* we first build a simplicial complex representation of 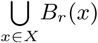. In this paper we focus on the Vietoris–Rips complex, but we note that there are several ways in which one can build a simplicial complex representation of 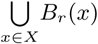.

We define a k-simplex as the convex hull of k + 1 affinely independent points, i.e. the k-simplex of *k* +1 affinely independent points is the convex polygon whose vertices are precisely the *k* +1 affinely independent points (21, 22). For example, a 0 simplex is a point, a 1 simplex is an edge, and 2 simplex is a triangle, and so on. Denote a k-simplex corresponding to the convex hull of 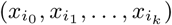 as 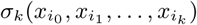. Then, the Vietoris–Rips complex of X with respect to *r* is the union of all k-simplices 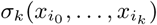 such that 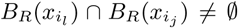 for all *l, j* = 0, 1, …, *k*. Note, another common simplicial complex is the Cech complex (mentioned in *Coalescent Intuition for Topological Statistics*), which instead requires non-empty mutual intersections of the k-simplices rather than non-empty pair-wise intersection.

The homology is computed on the simplicial complex representation of 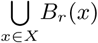 instead of the original union. Simplicial complexes are easier to work with computationally, and there exist theoretical guarantees that make it feasible to compute the homology of simplicial complex instead of 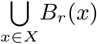 (See *Nerve theorem* in (21)).

Lastly, we take a sequence of parameters 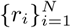, build the simplicial complex of X with respect to *r*_*i*_ and compute its homology for all *i*. This yields a sequence of a families of vector spaces associated to X, known as the persistent homology of *X*.

We represent the persistent homology of X with a barcode diagram ℬ_*i*_ for each dimension *i* (28). A bar (*b, d*) ∈ *ℬ*_*i*_ represents a generator of homology in dimension *i*. The birth time b of the bar corresponds to the *r* value at which the homological feature first appeared and the death time d of the bar corresponds to the *r* value at which the homological feature collapsed. Bars with longer bar length (d-b) are of particular interest since they persist throughout the sequence of simplicial complexes.

In this work, *X* is a collection of genomes and we use the Hamming distance *d*_*H*_ to build Vietoris Rips representations of sampled populations using *B*_*r*_(*x*) = {*y* : *D*_*H*_(*x, y*) ≤ *r*}. We then compute the persistent homology of *X* and extract statistics from the corresponding dimension-0 and dimension-1 barcode diagrams as input for the Topological Recombination Estimator.

### Filtering β_1_

Some authors on the subject of TDA have suggested that some bars in a persistence diagram may arise due to topological noise, especially those bars are that short-lived. It is unclear whether a universal threshold exists to filter out potentially noisy bars, however. In the case of our own study, we took it upon ourselves to test whether filtering out short-lived bars improves the relationship between existing topological models of recombination and *β*_1_, specifically using Camara et al’s *ρ*_*ph*_ for goodness of fit. We find that filtration of shortlived bars only lowers the *R*^2^ value for goodness of fit to this model in our simulated datasets, thus we opt not to use any filtration of these bars in practical applications (Figure S1).

### Missing Data

As large quantities of missing data are frequently encountered in empirical datasets, we investigated the performance of both *ψ* and *β*_1_ under various scenarios involving different amounts of missing data using the same datasets we simulated for our main coalescent investigation. Specifically, we took these existing datasets, duplicated them, and then converted randomly chosen sites or blocks of sites to N, in order to simulate missing data. For each dataset, a total of 10% of each sequence (and therefore, 10% of the total alignment) was converted to Ns. Since in all cases, 10% missing data was introduced into the alignments, we note that our specific interest here is in how robust each topological feature is to either a) a random distribution of missing sites, as would be common with sequencing error, or b) tandemly linked missing sites, as common in indels.

**Fig. 9.**
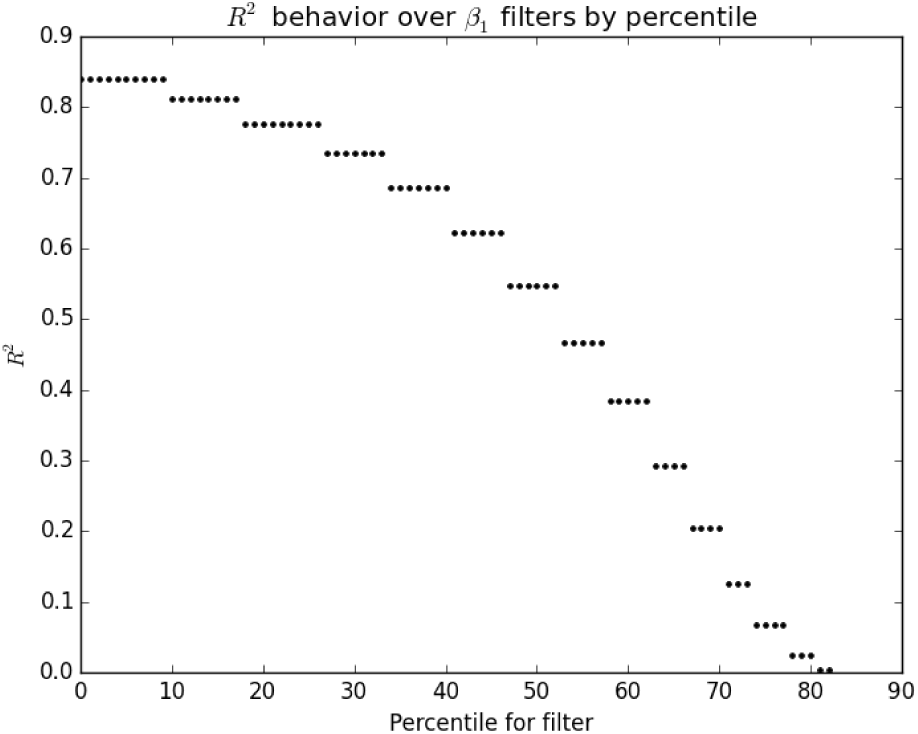
*R*^2^ values decrease as we filter out bars of increasing lengths in terms of percentiles of the overall distribution of bar lengths across all simulated datasets.

We find that, overall, *ψ* is much more robust to missing data than is *β*_1_ when predicting *ρ*. Specifically, we looked at the expectation of *β*_1_, *ψ* given *ρ* as in Camara et. al’s model, looking to see how well the model fits each topological feature under each missing data scenario. We did this in order to evaluate the strength of the relationship without a prior expectation for the expectation of *ψ*, and found a greater fit to the Camara model describing the expectation of *ψ* in place of *β*_1_ given *ρ* in these circumstances than their original expectation of *β*_1_ given *ρ*.

Looking at the reverse case, the prediction of *ρ* from *ψ* or *β*_1_ with missing data, we lose the fit to the Camara model, but may still observe differences in variance of the predictions and a clear relationship between *ψ* and *ρ* that is lost in *β*_1_ under these circumstances.

### Mixing Populations

We explored the robustness of *ψ* and *β*_1_ under mixed populations of varying recombination rate. In particular, we randomly sampled *N* individuals from a population of known recombination rate *ρ*_1_, along with *M* individuals from a population of known recombination rate *ρ*_2_. We kept *N* + *M* = 160 constant while varying N, M, *ρ*_1_, and *ρ*_2_. These experiments were all done on the simulated data, and we fixed *θ* = 25.

For each randomly concatenated population we computed the weighted mean recombination rate 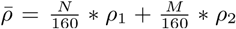. The results of comparing our main topological summary statistics *ψ* and *β*_1_ to the weighted mean recombination rate are presented in figures 12-13. We see that under randomly mixed populations *ψ* maintains a tight exponential relationship with 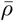. In comparison, in this setting the relationship between *β*_1_ and 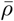 becomes noisier. The nice behavior of *ψ* is expected as mixing two distant populations only adds one non-informative coalescence event between the populations, and as a result has little effect on *ψ* other than averaging the recombination parameters of the samples.

**Fig. 10.**
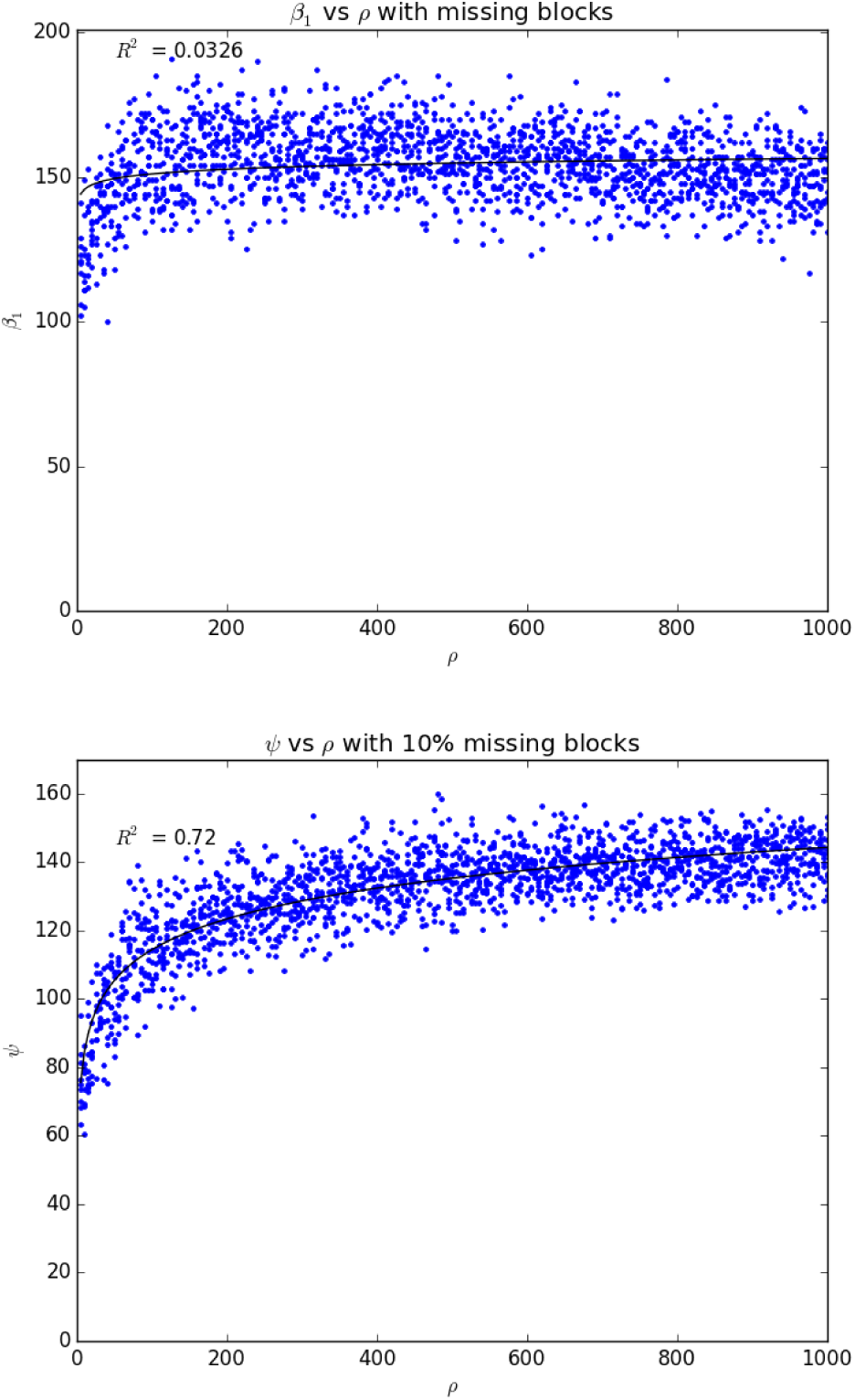
Expectations of *ψ* and *β*_1_ given 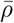 when 10% of data is missing in large tandem blocks. In the case of *β*_1_, the *R*^2^ value has dropped from 0.9 in the case of no missing data to 0.03, suggesting that this feature is highly sensitive to minimal missing data. In comparison, *ψ* maintains an *R*^2^ of 0.76, suggesting greater robustness to loss of information in sequence data.

### Lasso Weights

In order to gain intuition for which barcode statistics would be the best predictors for recombination we first ran a LASSO regression analysis using 15 barcode statistics as input and analyzed the outputted weight vector. We used the LASSO weight vectors as a proxy for the predictive power of each barcode statistic.

The barcode statistics we studied were the Betti number (*β*_*i*_), mean barcode length (*ψ*_*i*_), medium barcode length (*m*_*i*_), maximum barcode length (*M*_*i*_), and the thresholded Betti number 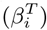 in dimensions 0,1, and 2. Here, the thresholded Betti number refers to the number of bars whose bar length is greater than a specified cutoff, where the cutoff is set as a percentage of maximum bar length *M*_*i*_. In these experiments we tested thresholds corresponding to 10 – 60% of the maximum bar length.

**Fig. 11.**
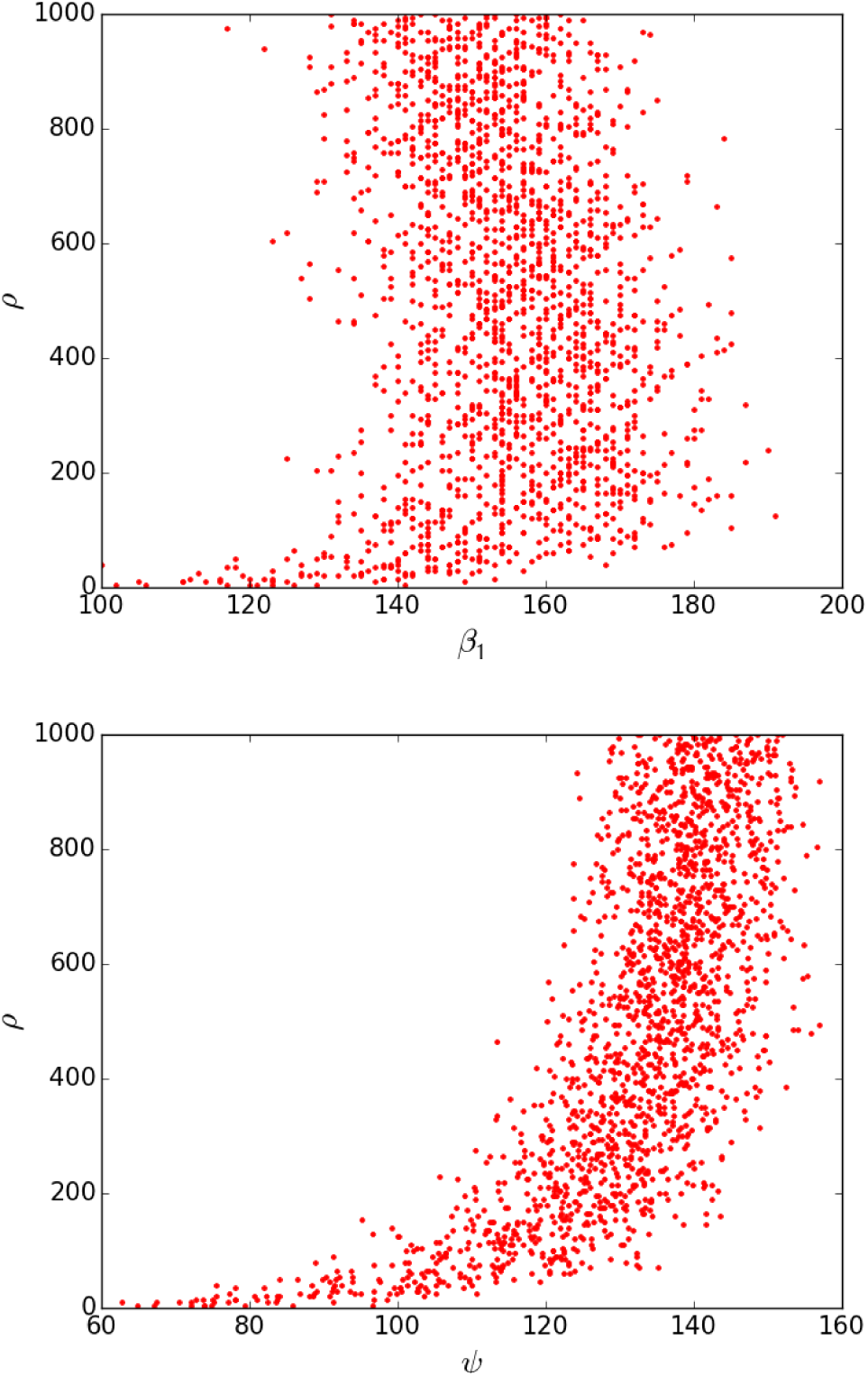
*β*_1_ and *ψ* as predictors of 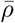 when 10% of data is missing in large tandem blocks. Though we lose the fit to the expectation model, we find a noticeable difference in the variance of the values of *ψ* and a visually tighter correlation to *ρ* than we see for *β*_1_.

We ran LASSO on the simulated data with fixed sample size using Scikit-learn (39) in Python 2.7. For each threshold, we ran LASSO 20 times on randomly selected training data. Initially we used all 15 barcode summary statistics as input and then analyzed the absolute vector of the LASSO weight vectors for all 20 runs across the varying thresholds. These weight vectors are visualized in a heat map in Figure 14.

Observe, consistently *ψ*_0_ (or *ψ*), *m*_0_, and 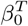 are among most heavily weighted inputs, with *ψ*_0_ having the most influence overall. This motivated our rigorous exploration of *ψ* as an estimator for recombination rate. Also note that *β*_0_ has zero weight since *β*_0_ is precisely the sample size, which is constant in this experiment. Consequently, 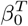 only varies as *M*_0_ varies.

In contrast to the dimension-0 statistics, the dimension-1 barcode statistics have negligible weights across all model runs and all thresholds. Moreover, almost all of the dimension-2 barcode statistics have low weights across all model runs, with the exception of *M*_2_ whose weight increases as the threshold for 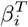 increases.

**Fig. 12.**
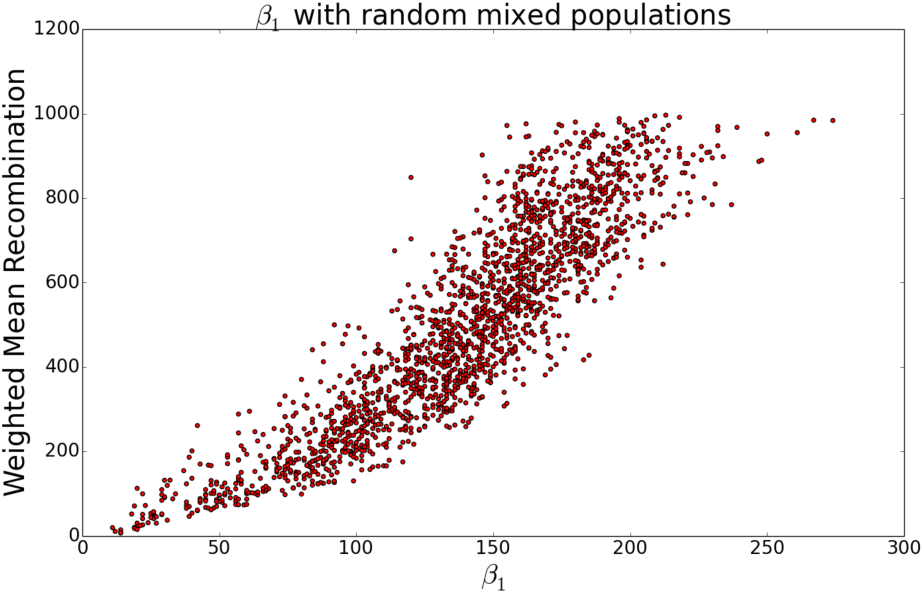
*β*_1_ versus 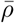 rate from randomly mixed populations of different recombination rates.

**Fig. 13.**
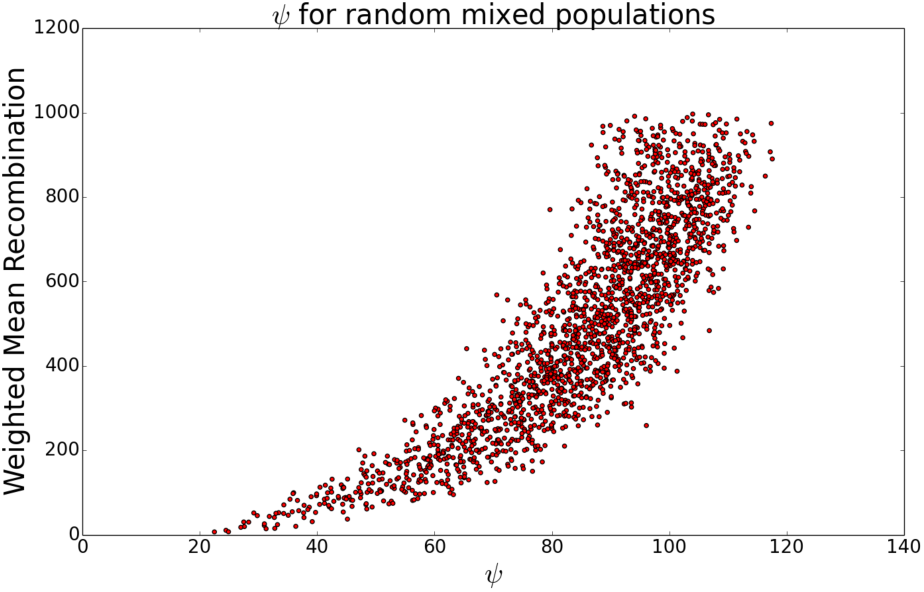
*ψ* versus 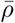 from randomly mixed populations of different recombination rates.

In an attempt to extract the best dimension-1 topological predictor for recombination, we re-ran the 20 runs of LASSO for varying thresholds using only the dimension-1 topological summary statistics as input. The results of this refined analysis are presented in Figure 15. Here we see that out of all the dimension 1 topological statistics, *β*_1_ is the most heavily weighted input feature when the threshold is set to ≤ 30% of *M*_*i*_. For increased threshold values the weight of *β*_1_ decreases significantly and *ψ*_1_ is the most heavily weighted input features for the dimension 1 barcode statistics, although its weights vary greatly across different model runs. This is consistent with the results presented in *Filtering β*_1_, which suggest that filtering out short-lived bars hinders the predictive power of *β*_1_ for recombination estimates.

The results of these LASSO analyses provided us with the motivation to focus on understanding the significance of lower dimensional topological statistics as predictors for recombination. Moreover, we used the results of the dimension-1 LASSO analysis to decide which higher dimensional statistics to use in tandem with *ψ*. We chose to exclude dimension-2 statistics as predictors due to their overall low weight vectors and the lack of biological significance.

**Fig. 14.**
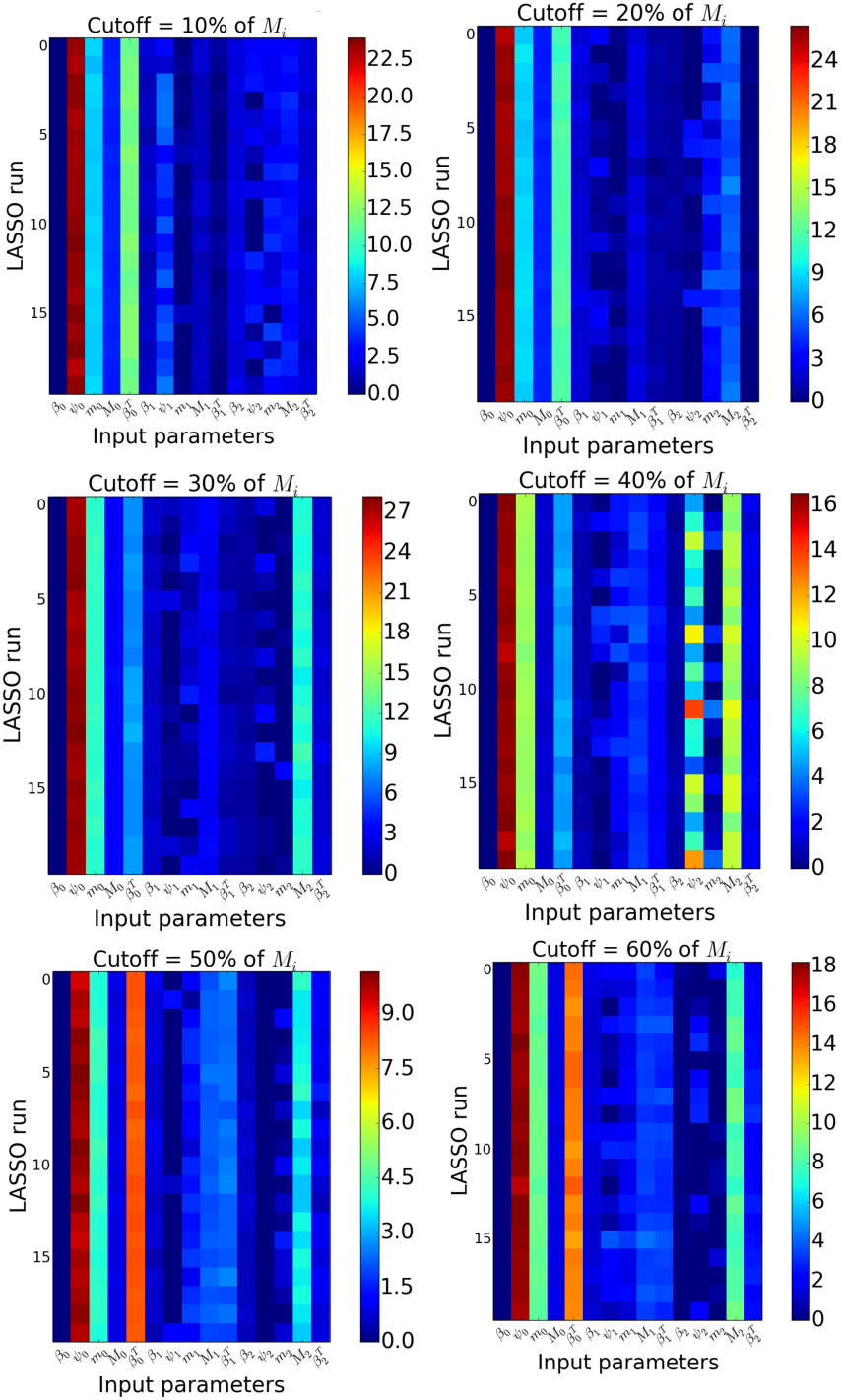
Absolute value of LASSO weight vectors across 20 model runs on randomly selected 15-dimensional training data for varying thresholds for 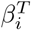.

### Noise Experiments

We further tested whether topological noise may contribute to relationships we observed between *β*_1_, *ψ*, and *ρ* by adding noise or randomizing sequences within datasets to obtain topological structures unrelated to recombination. In one experiment, we simulated realistic cases of sequencing error in each dataset by adding a random base over a range of error rates from 0 to 140 / 1000 bases. As this error rate increases, we find a reduction in the fit of topological models in *β*_1_ to the data, but the decrease in fit is slow over all realistically expected error rates (around 10% sequencing error, we see no reduction in *R*^2^ for *ρ*_*ph*_.

### Other Relationships

We tested for relationships between our topological features *ψ* and *β*_1_ and Watterson’s *θ*, as well as for possible correlations between the topological features themselves.

We computed each feature from the set of 3600 simulated sequence alignments with varying values of *ρ* and *θ*, and produced similar correlation plots showing relationships between each pair of variables. We find that neither *β*_1_ nor *ψ* is correlated with Watterson’s *θ*, thus we can confidently assert that this is not a confounding factor in our analyses (Figure 17).

In contrast to this, we do find that *β*_1_ and *ψ* are correlated quite tightly in our work (Figure 18). Though this is the case, we have noted in other experiments that *ψ* nevertheless adds unique information.

**Fig. 15.**
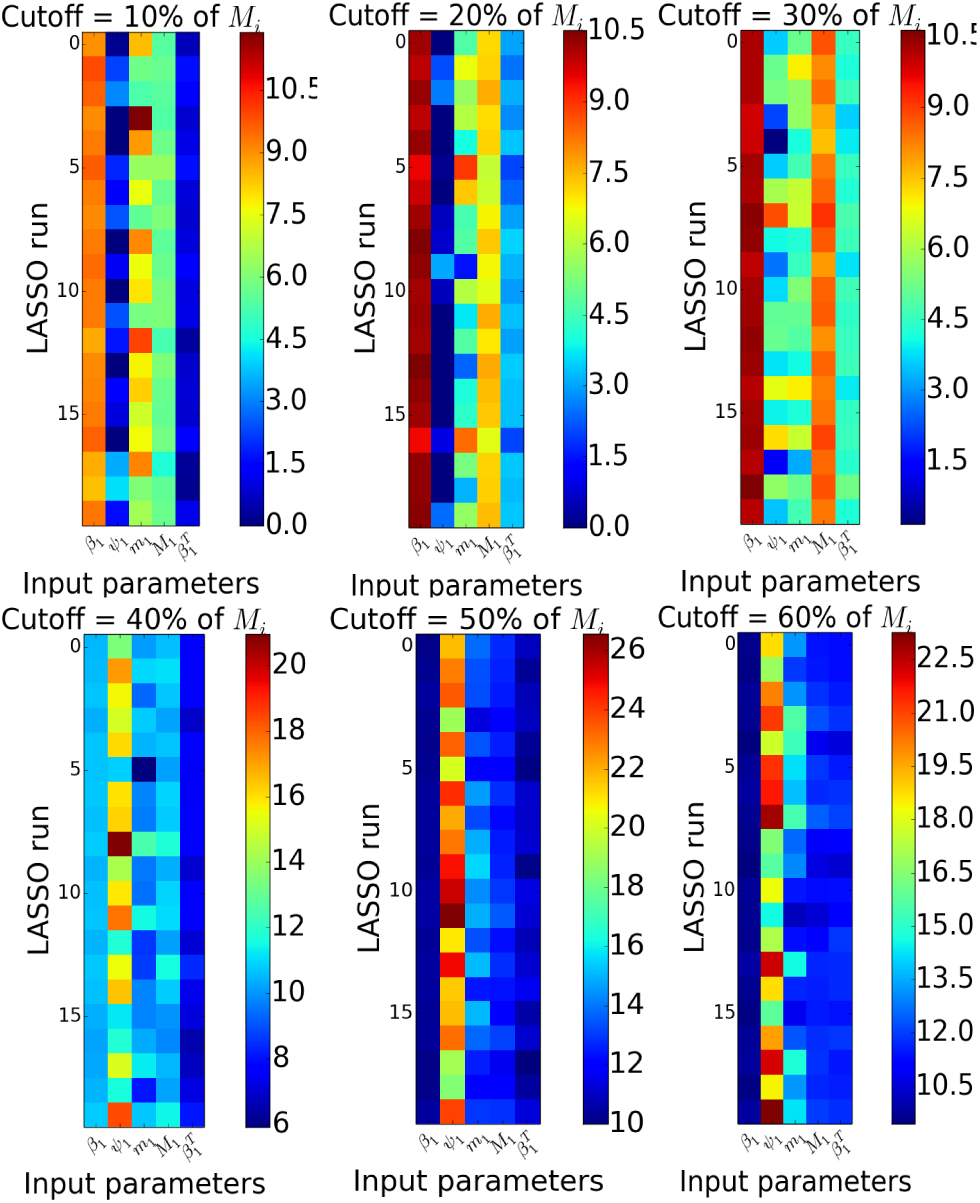
Absolute value of LASSO weight vectors across 20 model runs on randomly selected dimension 1 training data for varying thresholds for 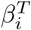.

**Fig. 16.**
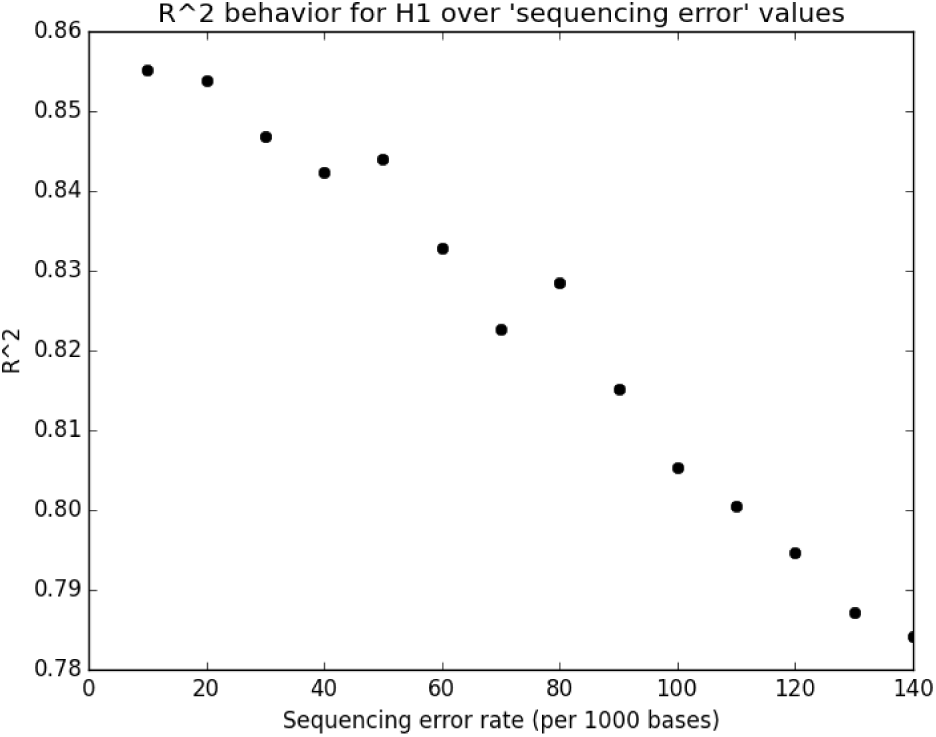
*R*^2^ values decrease as we include increasing sequence errors, but remains greater than 0.75 over all realistic and slightly more extreme possible error rates.

**Fig. 17.**
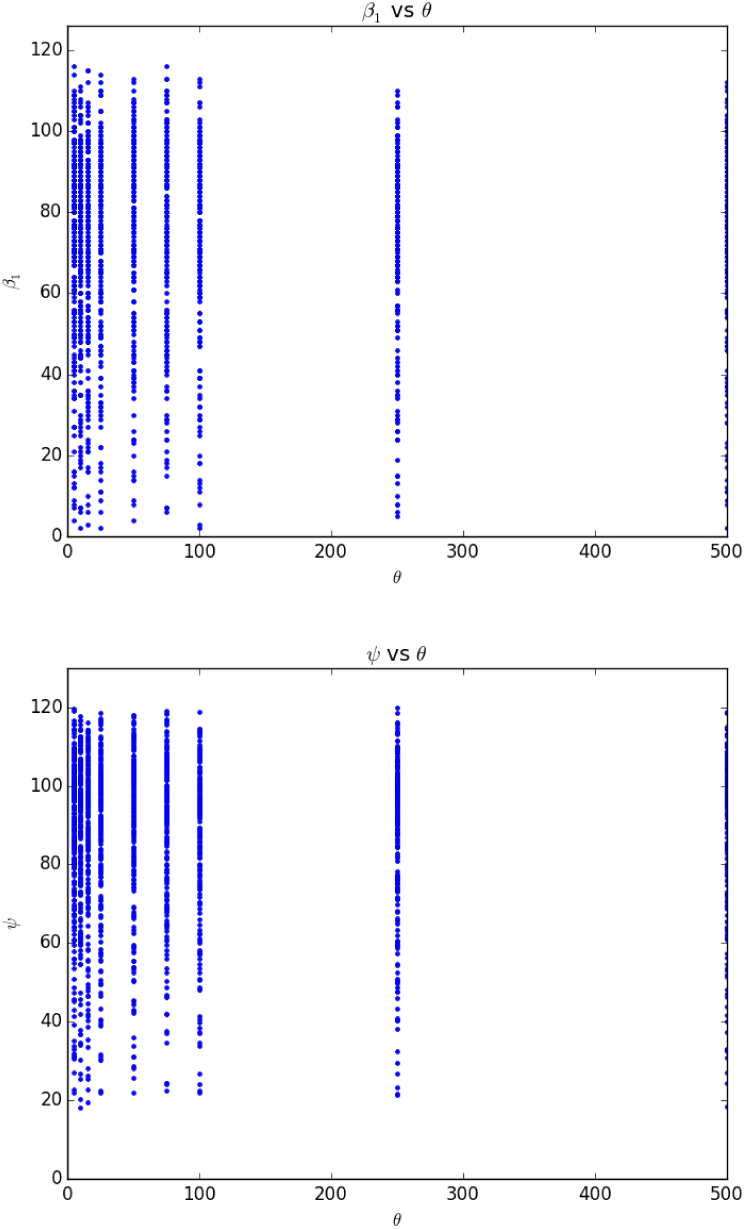
*ψ* and *β*_1_ are uncorrelated with variance in *θ*.

**Fig. 18.**
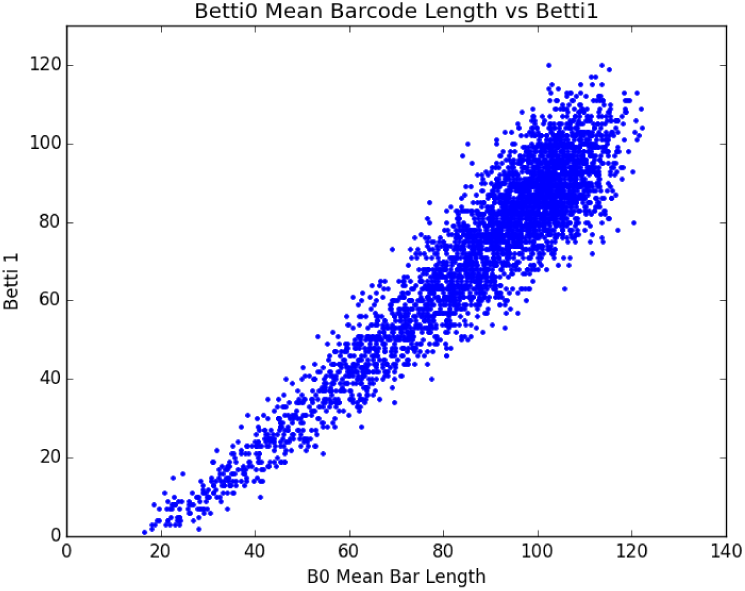
*ψ* and *β*_1_ are tightly correlated over variable values of *ρ*.

Author contributions
D.P.H., M.R.M, M.M., and A.J.B. designed and conducted experiments;D.P.H., M.R.M., and M.M. wrote code for the project; D.P.H. and M.R.M performed empirical analyses, M.M. contributed theory; and D.P.H., M.R.M, M.M.,and A.J.B. wrote the paper

